# Quantifying the Effect of Encapsulation on Microbial Mineralization Processes Using a Modified DNA Extraction Method

**DOI:** 10.64898/2026.10.05.756701

**Authors:** Eva M. Albalghiti, Jeremy D. Semrau, Brian R. Ellis

## Abstract

Microbially-mediated mineral precipitation is important in the natural world and to a number of emerging climate, energy, and water technologies. Central to many microbial mineralization processes is the preferential accumulation of precipitates on microbial surfaces. In some systems, including ureolytic microbially-induced carbonate precipitation (MICP), precipitates may entomb cells in a process termed “encapsulation.” Encapsulation is known to greatly affect microbial systems, but understanding these relationships quantitatively has proven difficult, largely because nucleic acid extraction from encapsulated cells yields inconsistent results.

Here, a demineralization protocol adapted from studies of natural microbialites is shown to recover genomic DNA from carbonate-encapsulated *Sporosarcina pasteurii* at yields indistinguishable from unmineralized cells and of sufficient quality for quantitative PCR. This method allows the degree of encapsulation to be expressed quantitatively as the amount of precipitate formed normalized to the estimated number of cells present, and allows growing populations to be tracked with increased precision. Simultaneous growth and mineralization experiments in mixed batch reactors suggest encapsulation-induced growth suppression at calcium concentrations of 250 mM, which are commonly employed in MICP under different physicochemical conditions. Variable rates of precipitate accumulation on cells, possibly stemming from local mass transport constraints, are proposed as a mechanism of calcium concentration-dependent growth suppression. Accordingly, mass transport principles are used to derive a reaction rate scaling function, allowing the observed growth and precipitation behavior to be replicated by a numerical model. These findings lay the groundwork for a more fundamental understanding of how encapsulation shapes microbial mineralization processes.

**Importance:** Some microorganisms have the ability to form minerals from dissolved ions, which often build up on microbial cells and biofilms. We use this phenomenon to answer key questions in geology and ecosystem science, and to engineer new solutions to environmental problems. When microorganisms become fully encased in precipitates, however, it can pose problems for extracting microbial DNA, which limits our biological understanding of these systems. In this study, we use a model calcium carbonate-precipitating bacterium to test a modified DNA extraction approach. This approach allows us to obtain high and reproducible yields of good-quality DNA even from completely encapsulated cells. We then show how this method can help us better understand the fundamental relationship between carbonate precipitation and cell growth.

## 1. Introduction

Microbially-mediated mineral precipitation processes are known to be both highly influential in the natural world^1^ and of great potential utility in engineered systems^2^. Among the microbial mineralization processes targeted for engineering uses, ureolytic microbially-induced carbonate precipitation (MICP) is perhaps the best-studied example^3–5^, with a growing list of environmentally-relevant applications that includes self-healing concrete^6,7^ as well as subsurface engineering applications such as groundwater remediation^8,9^ and enhanced geologic carbon sequestration^10–13^. In subsurface applications, the tendency of microbes to act as mineral scaffolds^14^ is essential. While the same carbonate precipitation pathway could be easily mediated using, for example, purified urease^15–17^, the preferential nucleation and growth of precipitates on microbial biomass^18,19^ allows for greater control over the timing and distribution of precipitate formation^20,21^, provided that microbial attachment and transport behavior is well-characterized^22,23^.

The accumulation of carbonate precipitates on microbial surfaces has been shown to lead to encapsulation^24^, in which microbial biomass becomes fully entombed within precipitates^14^. It has been suggested that encapsulation affects the growth and ureolytic activity of microorganisms, thereby shaping the quantity, distribution, timing, and efficiency of precipitate formation^24–26^. Many MICP models incorporate encapsulation by designating an encapsulation-specific cell decay coefficient, which can be a function of the precipitation rate or the fraction of the domain not currently occupied by precipitates^11,27,28^. Alternative approaches include relating the effects of encapsulation to the ratio of an assumed maximum rate of carbonate precipitation and a theoretical encapsulation rate constant^29^, or integrating the hypothesized effect of encapsulation on cell viability into other processes without modeling encapsulation specifically^30,31^. All of the above approaches rely on parameters that are difficult to constrain experimentally, and it has been shown that modeled outputs are sensitive to these parameters^29^. This suggests a need for quantitative characterization of the mass transport and kinetic constraints on encapsulation, and its effect on cell growth, under varied experimental conditions.

Despite the acknowledged importance of encapsulation, there has been little quantitative exploration of how encapsulation affects microbial growth and precipitate accumulation. A salient barrier is the limited and inconsistent yields of genomic DNA extracted from carbonate-encapsulated cells^32^. Without a robust DNA extraction methodology, total cell numbers cannot be reliably estimated, rendering systems that involve simultaneous growth and mineralization difficult to resolve. Issues with DNA extraction can also introduce uncertainty in analyses of carbonate-precipitating microbial communities, an essential tool in evaluating biostimulated MICP approaches^9,33^.

Outside of the engineered MICP literature, research on natural microbialites has produced diverse methods for extracting DNA from carbonate-containing samples, many of which involve a demineralization pre-treatment step prior to DNA extraction. While some studies have reported the successful use of acidic solutions for demineralization^34^, others note that low pH leads to extensive DNA degradation in many sample types^35^. An alternative demineralization method is chelation using alkaline ethylenediaminetetraacetic acid (EDTA)^35,36^, which has previously been used for DNA extraction from mineralized tissues such as bones and teeth^37^, and which is a common component of nuclease-inhibiting DNA storage buffers. The ability of chelators to inhibit polymerase chain reaction (PCR) enzymes is, however, perceived as a barrier^38^.

In this study, a modified method for DNA extraction from carbonate precipitates is demonstrated on encapsulated *Sporosarcina pasteurii* cells. The proposed method is shown to reproducibly promote previously-unattainable DNA yields, and its efficacy is found to depend on the extraction of DNA from both the demineralization supernatant and the demineralized cell debris. The method is then applied to analyzing mineralization processes in mixed batch reactor experiments, using both inactive and growing cell populations. Experiments on inactivated cells are used to estimate the maximum amount of precipitate that can be expected to accumulate per intact cell under these experimental conditions, and to identify sources of variability in this estimate. Simultaneous growth/mineralization experiments show that high calcium concentrations can inhibit cell growth independent of any ionic strength effect. Calcium concentration-dependent growth suppression is hypothesized to stem from calcium concentration-dependent differences in maximum precipitation rates, which allow encapsulation to outpace cell growth at high concentrations. The proposed relationship between calcium concentration, encapsulation rate, and cell growth is further explored through simplified numerical models, where experimental results are successfully reproduced using a precipitation rate scaling function derived from independently-sourced mass transport parameters. Taken together, these results lay an essential foundation for mechanistic investigations of encapsulation and its effect on cell growth and precipitate yield in microbial mineralization systems.

## 2. Results and Discussion

### 2.1 Demonstration Trials Using Inactive *S. pasteurii*

Figures 1-3 summarize results from the three growth-free demonstration trials. These trials served two purposes. The first was to evaluate the efficacy of the applied demineralization procedure for facilitating genomic DNA extraction from carbonate-encapsulated *S. pasteurii* cells, and to identify potential sources of variability in DNA yields (Figures 1-2). The second was to derive a realistic upper bound on per-cell precipitate formation in the absence of time and dissolved calcium limitation, and to identify potential sources of variability in per-cell precipitation (Figure 3). Accompanying numerical data on “no mineralization” control experiments, including colony count assay results used for interconversion between gene copies, OD600, and cfu/mL, can be found in Table S1.

**Figure 1.**
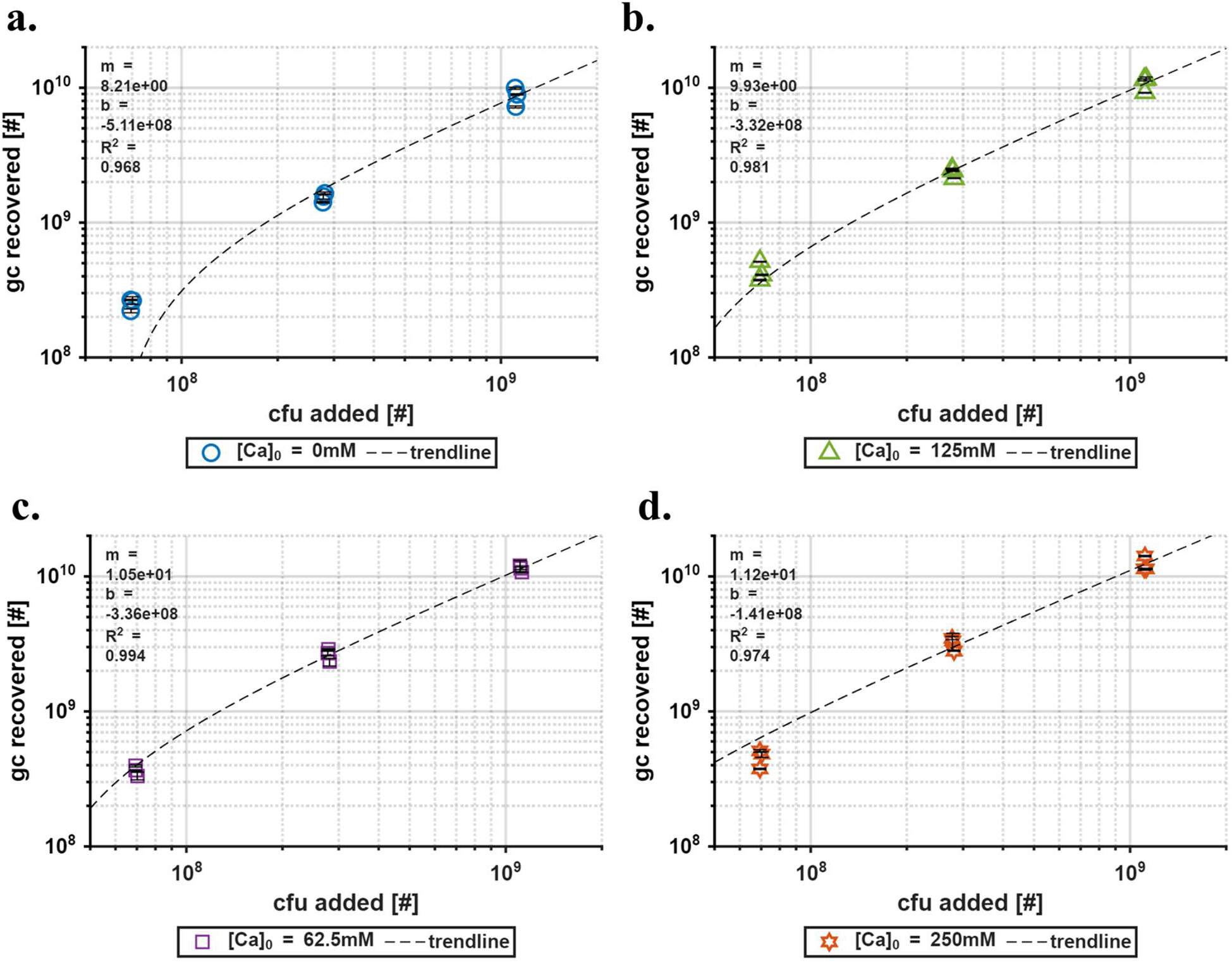
Number of gene copies recovered in demonstration trials *versus* number of cfu added, across three trials and three cell densities, for samples with initial calcium concentrations of a.) 0 mM, b.) 62.5 mM, c.) 125 mM, and d.) 250 mM.

**Figure 2.**
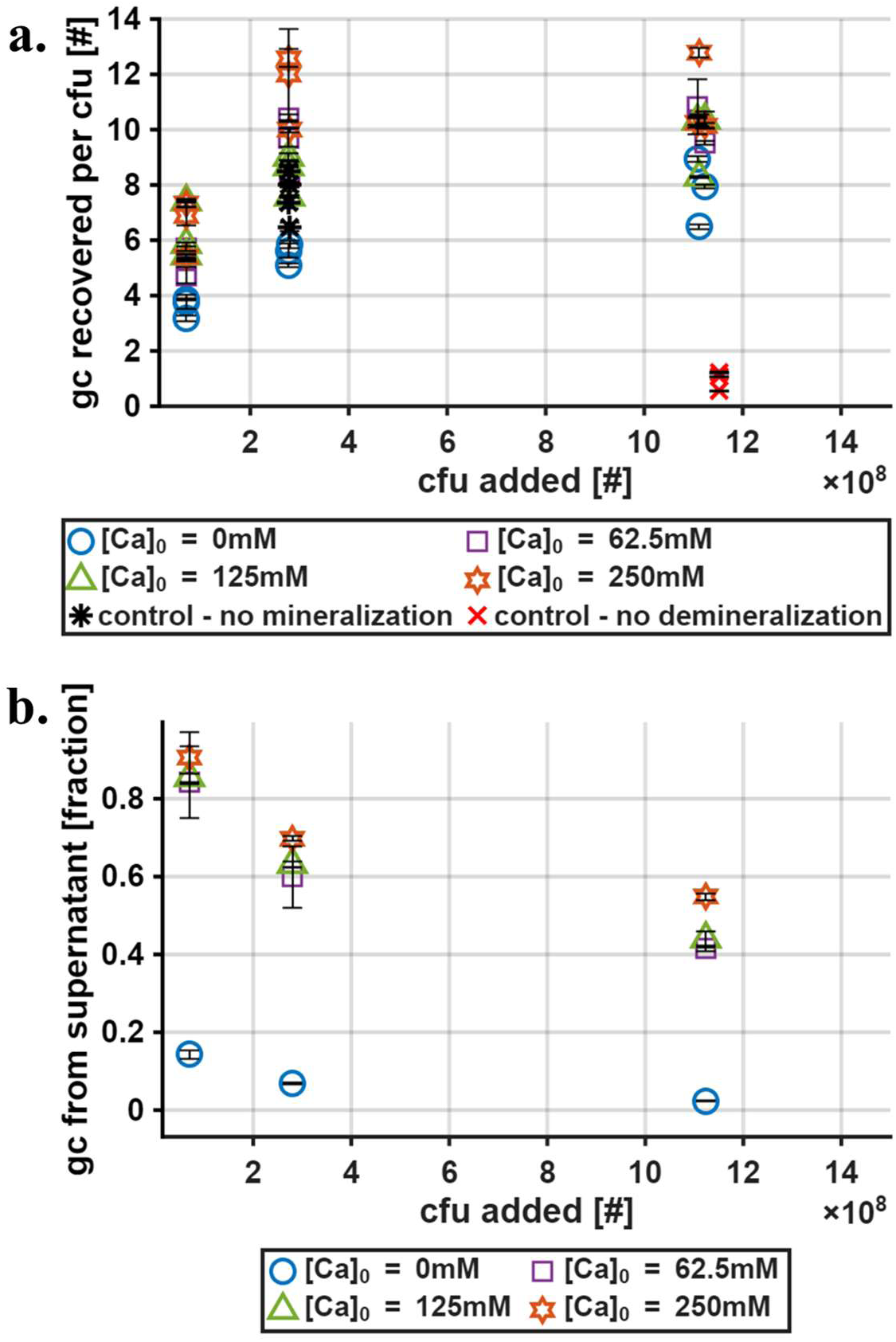
a.) Number of gene copies recovered per cfu added *versus* total number of cfu added for four initial calcium concentrations, alongside “no mineralization” and “no demineralization” control samples. b.) Fraction of gene copies recovered from the supernatant concentration portion of the extraction, across four calcium concentrations and three cell levels.

**Figure 3.**
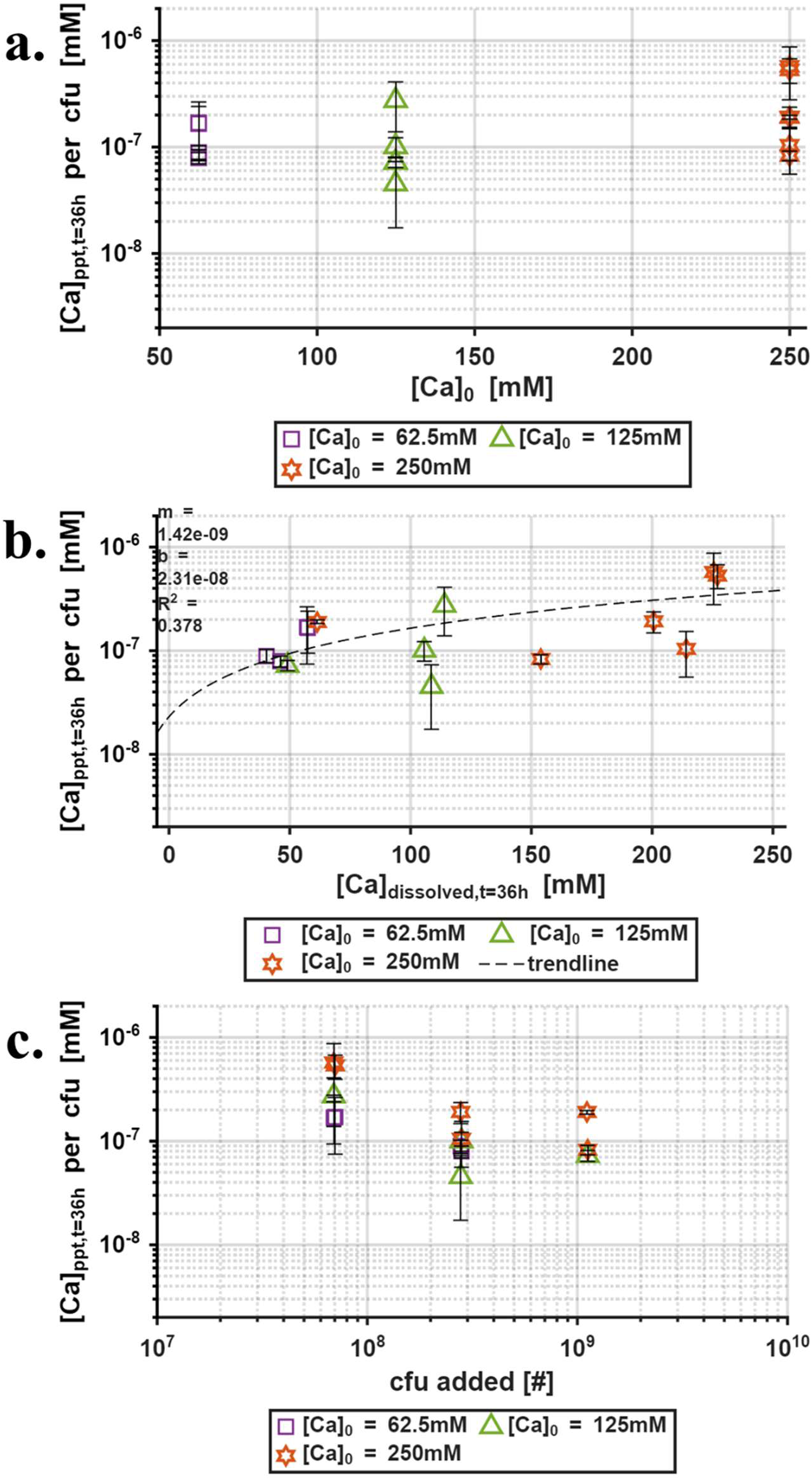
Average amount of calcium carbonate precipitate accumulated per cfu added over the course of 36 h demonstration trials, *versus* a.) initial calcium concentration, b.) final calcium concentration, and c.) total number of cfu added.

#### 2.1.1 DNA Extraction Method Evaluation

Figure 1 demonstrates that the number of gene copies recovered has a robust, linear dependence (R^2^ range 0.968-0.994) on the number of cfu added for cfu loadings between 7.0x10^7^ and 1.1x10^9^, irrespective of initial calcium concentration. In other words, DNA yields are consistent across a range of cell densities and mineralization conditions. Trendline y-intercepts range from -5.1x10^8^ and -1.4x10^8^ gc, while slopes range from 8.2 to 11.2 gc/cfu, with steeper slopes and less negative intercepts corresponding to higher initial calcium concentrations. Since *S. pasteurii* is known to have eight 16S rRNA copies^39^, a y-intercept of 0 gc and a slope of 8 gc/cfu would be expected with perfect DNA recovery and a 1:1 correspondence of cfus to cells. Negative intercepts therefore imply a lower bound on the number of cells from which DNA can be reliably extracted, while slopes greater than 8 imply that colony count assays underestimate the total number of cells. These are generic limitations of the DNA extraction and cell enumeration methods used here, and not specific to the tested demineralization procedure. Limitations of these experimental methods notwithstanding, trendline slopes suggest high and consistent DNA yields across all initial calcium concentrations.

Figure 2a further contextualizes the above findings by comparing DNA yields to those observed in both “no mineralization” and “no demineralization” control experiments. Note that the difference between “[Ca]_0_ = 0 mM” and “control - no mineralization” is that the former received otherwise identical treatment to 62.5, 125, and 250 mM samples, while the latter was neither diluted into a mineralization reaction tube nor subjected to demineralization treatment (section 3.2.2). No statistically significant difference in DNA yield was found between the “no mineralization” control and mineralized samples of any initial calcium concentration. In contrast, DNA yields from demineralized samples appear to exceed those from the “no demineralization” control by up to an order of magnitude, a statistically significant difference (p-value: 0.003). These results suggest that the demineralization treatment successfully promoted high DNA yields from previously encapsulated cells, comparable to yields from never-mineralized cells (i.e. “no mineralization” control).

It is important to note that while there is some variability inherent to DNA extractions from even never-mineralized cells (standard deviation of “no mineralization” control yields: 1.25 gc/cfu), there is greater variability among samples subjected to mineralization and demineralization (pooled standard deviation: 6.19 gc/cfu). Furthermore, DNA yield appears to have some dependence on both cfu loading and initial calcium concentration. As expected, lower yields are obtained with cfu loadings of 7.0x10^7^ compared to cfu loadings of 2.8x10^8^ and 1.1x10^9^ cfu, although this difference is not statistically significant. Interestingly, the effect of initial calcium concentration runs contrary to expectation, with higher calcium concentrations corresponding to higher yields. This finding aligns with the steeper trendline slopes observed with increasing calcium concentration in Figure 1.

The effects of cfu loading and initial calcium concentration are further explored in the supplementary information (Figures S1-S2). Briefly, these results suggest that the extraction method used here likely underestimates the number of cells present for cfu loadings less than 2.8x10^8^ total cfu, or 1.4x10^7^ cfu/mL. Similarly, the number of cells present may be overestimated at high calcium concentrations. The magnitudes of these effects are not, however, large enough to produce gc/cfu recovery rates that differ significantly from the unmineralized control, suggesting that estimates are still within a reasonable margin of uncertainty for DNA extraction procedures. Accordingly, when the number of gc recovered from a sample is subsequently used to estimate cfu_eq_ (section 3.6), a single conversion factor derived from the “no mineralization” control (7.79 gc/cfu, see Table S1) is applied regardless of hypothesized cell density or calcium concentration. The uncertainty embedded in this conversion is contained in the standard deviation (1.25 gc/cfu), which is propagated into the final reported value (section 3.7).

A potential reason for decreased yields at the lowest cfu loading may be incomplete recovery of microbial biomass from the 20 mL mineralization reactions. Here, separation of cells from both mineralization medium and demineralization solution was achieved through centrifugation at 4500 rcf for 30 min (section 3.2). Centrifugation at higher g values might therefore improve recovery rates. This may also partly explain increasing yields at higher calcium concentrations, since the accumulation of carbonate precipitates would improve pelleting by increasing the density of cells and aggregates. Another source of DNA loss might be the lysis of some cells in 0 mM reaction tubes resulting in DNA leakage to the mineralization medium, which is discarded. This effect might be mitigated in calcium-containing samples through protective encapsulation of biomass, and possibly sorption of leaked DNA to precipitates.

To further interrogate how the carbonate encapsulation process might contribute to elevated DNA yields, amounts of DNA extracted from cell pellets and concentrated from demineralization supernatants were compared (Figure 2b). Although some DNA was successfully recovered from both the pellet and supernatant in all samples, the relative proportions are found to vary significantly. In mineralized cells, between 40 and 90% of gc recovered were found to originate from the demineralization supernatant, compared to a maximum of 15% in unmineralized cells. Lower cell numbers and higher initial calcium concentrations generally correlate with a higher fraction of DNA recovered from the supernatant. This points to DNA leakage into the demineralization supernatant, with the extent of leakage directly related to the extent of cell encapsulation. Mineralization followed by demineralization treatment may therefore increase DNA yields by promoting more complete cell lysis. It should also be noted explicitly that achieving high and consistent yields appears to require that DNA be extracted from both the cell pellet and the demineralization supernatant, which is not currently standard practice in studies employing similar demineralization methods^35,38,40^.

#### 2.1.2 Estimating Maximum Per-Cell Precipitation

In addition to validating the efficacy of the applied demineralization method, the demonstration trials also served to establish bounds on expected per-cell precipitation for these experimental conditions. Figure 3 compares per-cell precipitation after 36 h of mineralization across the three cfu loadings and three initial calcium concentrations. At the same time, it was important to assess whether per-cell precipitation can be reliably derived from the number of gc recovered through cfu_eq_ (section 3.6), as this enables per-cell precipitation to be estimated even when the number of cfu present is not known *a priori*. Per-cell precipitation is therefore calculated using both the known cfu loading (Figure 3) and the number of gc recovered (Figure S3), converted to cfu_eq_ (section 3.6) using the conversion factor derived from unmineralized controls (Table S1). In order to remove calcium limitation as a confounding influence, this analysis excludes samples whose calcium concentration at 36 h was less than 10% of its initial value (section 3.6). The per-cell precipitate quantities shown here therefore represent an upper bound on the average, per-cell precipitate accumulation for these experimental conditions. As shown in Figure S4, removing this restriction skews the data in the negative-y direction as some samples fully deplete dissolved calcium before cells become fully encapsulated.

As Figure 3a shows, calculated per-cell precipitate quantities were found to range from 4.5x10^-8^ mM/cfu to 5.5x10^-7^ mM/cfu, a difference of approximately one order of magnitude. These are equivalent to 9.0x10^-13^ - 1.1x10^-11^ moles of calcium precipitated per cell, or 30 - 400 μm^3^ of calcium carbonate, assuming a typical molar volume of 36.9 cm^3^/mol. This is significantly larger than the volume of a typical *S. pasteurii* cell, estimated to be approximately 4μm^3^ assuming a cylindrical cell 5μm long and 1μm in diameter. Furthermore, no statistically significant differences in per-cell precipitation were identified based on initial calcium concentration. This suggests that as long as dissolved calcium remains in excess and time is not a limiting factor, the effect of initial calcium concentration on per-cell precipitate accumulation is minimal.

Sources of variability in per-cell precipitate accumulation are further explored in Figure 3b-c. The final calcium concentration (Figure 3b) was deemed potentially important because precipitation rate is expected to decrease as dissolved calcium concentration decreases. Whereas preliminary experiments used to determine the timescale of mineralization were conducted at the highest concentration of 250 mM (section 3.2 and supplementary information), samples in which dissolved calcium reaches significantly lower values might require longer times to reach full encapsulation. This hypothesis is consistent with the observed weakly positive correlation between final calcium concentration and per-cell precipitation (Figure 3b), for which the trendline slope was found to differ from zero to a statistically significant degree (p-value < 0.05). At the same time, low coefficient of determination (R^2^ < 0.4) indicates that this phenomenon can only explain a minority fraction of variation in per-cell precipitation.

The influence of cfu loading on per-cell precipitation was also examined (Figure 3c). Here, samples with the lowest cfu loading display visibly elevated per-cell precipitate quantities. A possible explanation for this effect may be that denser cell suspensions form more and larger aggregates or flocs, which could limit per-cell precipitation by restricting calcium transport to cells on floc interiors. While this difference was not found to be statistically significant, it does suggest that the effect of cell aggregation on mass transport will be important to a more complete understanding of encapsulation.

In summary, investigating per-cell precipitation in inactive cells yielded several insights that form a foundation for subsequent analyses. First, the theoretical maximum amount of precipitate that can be expected to accumulate per cell in these conditions is suspected to fall in the range of 9.0x10^-13^ - 1.1x10^-11^ moles of calcium. As these are the first empirically-derived estimates of precipitation normalized to cell numbers, they provide a useful benchmark against which to compare precipitation in actively growing cell populations, as well as an important parameter for mechanistic models.

Second, average per-cell precipitation is found to vary considerably across samples. While differences in initial calcium concentration had no apparent effect, it is shown that lower dissolved calcium concentrations near the end of mineralization are associated with less per-cell precipitation. Further resolving this relationship is beyond the scope of this study, but it is consistent with the hypothesized relationship between calcium concentration and precipitation rate. Similarly, the observation that lower cfu loadings produce higher per-cell precipitation suggest that probing microbial aggregation behavior during precipitation processes will be key to a more complete understanding of mass transport constraints during encapsulation.

Finally, a comparison between Figure 3 and Figure S3 demonstrates that cfu_eq_ (section 3.6), which represents inferred cell numbers based on DNA yields, reliably captures the same trends in per-cell precipitation as known cfu loadings. The conversion from DNA yields to cfu_eq_ introduces additional uncertainty (section 2.1.1, Figure S1-S2), which must be incorporated into analysis. For example, the effect of cfu loading on per-cell precipitation appears exaggerated in Figure S3c compared to Figure 3c, owing to the established underestimation of cell numbers at low cfu loadings (Figure 2a, Figure S1). There were not, however, any instances in which the conversion obscured or fundamentally changed underlying trends. The DNA extraction method applied here can therefore be applied to analyzing experiments on growing cells, where the true number of cfu present is unknown. This is done in subsequent analyses of simultaneous growth/mineralization experiments.

### 2.2 Simultaneous Growth/Mineralization Trials

#### 2.2.1 Effect of Calcium Concentration on Growth Rate

Achieving consistent DNA extraction from encapsulated cells allows for quantitative insight into the dynamics of simultaneous growth and mineralization. Figure 4 summarizes the results of three simultaneous growth and mineralization trials, showing the estimated number of cells present in cultures grown for time periods up to 36 h (Figure 4a, c, e), and the concentration of dissolved calcium measured in these cultures at the time of growth stoppage (Figure 4b, d, f). For added context, Figure S5 summarizes pH and dissolved oxygen data from growth/mineralization trials, and Figure S6 shows the results of two control trials, one conducted in lower-ionic strength growth medium and the other conducted without the 180 rpm mixing used in other trials (section 3.2). It is also important to note that the comparability of cfu_eq_ between inactive and growing populations relies on the as-yet untested assumption that the number of genome copies in *S. pasteurii* cells does not vary with growth phase, as has been observed in other organisms^41,42^.

**Figure 4.**
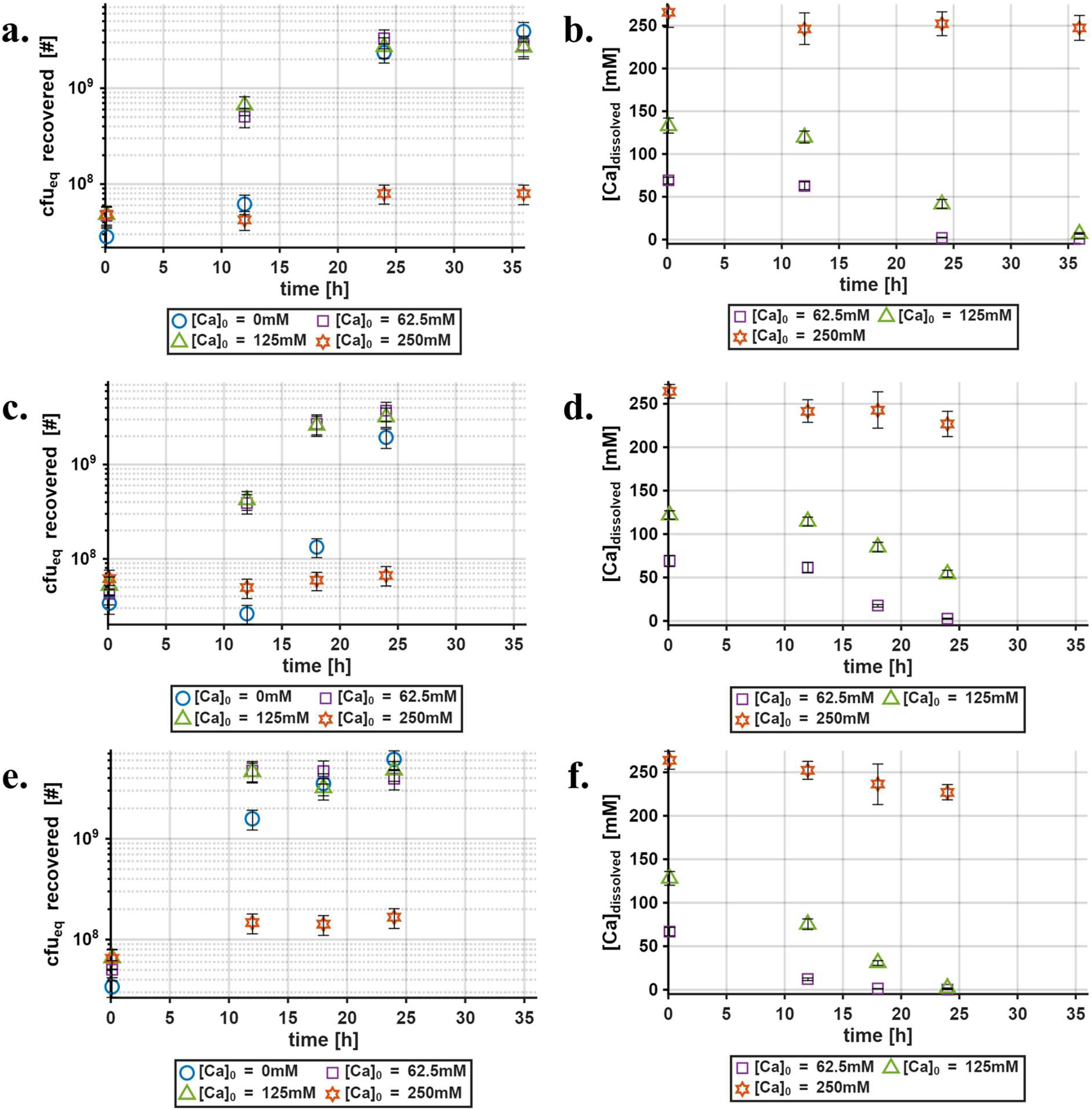
Summary of inferred cell growth (a, c, e) and calcium precipitation activity (b, d, f) across three simultaneous growth/mineralization trials (Trial 1 = a, b; Trial 2 = c, d; Trial 3 = e, f).

Based on inferred cell growth trends (Figure 4a, c, e), low-to-moderate concentrations of calcium (62.5 and 125 mM) do not appear to have a significant detrimental effect on *S. pasteurii* growth, as estimated cell numbers at 24 h are comparable across initial calcium concentrations of 0 mM, 62.5 mM, and 125 mM. Strikingly, almost negligible growth is observed at initial calcium concentrations of 250 mM, a difference found to be statistically significant at both 12 h (p-value: 0.016) and 24 h (p-value: 0.0001). Growth suppression at 250 mM of dissolved calcium differs from conventional wisdom derived from flowthrough experiments. For example, a recent study by these authors confirmed the growth of *S. pasteurii* in columns at dissolved calcium concentrations up to 300 mM^43^, and concentrations in this range and higher are frequently employed in the MICP literature^3,31,44^. This discrepancy may stem from mixing and mass transport limitations constraining the rate of precipitate accumulation on cells, as the spatial heterogeneity of flowthrough systems are expected to create calcium-depleted microenvironments not prevalent in the mixed batch system studied here.

Tracking dissolved calcium concentration over time (Figure 4b, d, f) adds important context in two ways. First, it is observed that dissolved calcium concentrations remain significant throughout periods of exponential growth, best illustrated by the persistence of dissolved calcium through 24 h in samples with initial calcium concentrations of 125 mM. This confirms that there is a range of dissolved calcium concentrations in which growth and mineralization co-occur, and growth is not simply initiating after calcium is depleted. Second, an analysis of changes in dissolved calcium concentration during each time interval suggests that precipitation proceeds more slowly here than would be predicted from whole cell *S. pasteurii* ureolysis kinetics alone^45^. Again considering samples with 125 mM initial calcium concentration as an example, estimated total cell numbers exceed 2x10^9^ cfu_eq_ by 18 h in trial 2 and 12 h in trial 3 (Figure 9b-c). Based on the per-cell precipitate accumulation of 10^-7^ mM/cfu observed at this cfu level and calcium concentration (Figure 8c-d), up to 200 mM of dissolved calcium could theoretically be precipitated by this amount of biomass even if no further growth occurred. Even at conservative estimates of urea concentration as low as 20% of the initial 300 mM concentration (section 3.1), accounting for only known kinetic constraints on ureolysis activity would predict a decrease in dissolved calcium of 100 mM in 1 h. Although progressive slowing of ureolysis activity is expected as encapsulation proceeds, this estimate stands in contrast to the less than 50 mM of dissolved calcium depletion actually observed over the following 6 h interval in trials 2 and 3. An analysis of trial 1 data for the period of 12 to 24 h yields a similar discrepancy. These discrepancies indicate that even in the mixed batch system used here, precipitation may occur at only a fraction of its theoretical rate even before significant encapsulation occurs, and precipitate accumulation may lag behind cell growth to a greater extent than would be expected in the absence of other limitations. The relationship of this phenomenon to calcium concentration and its potential influence on cell growth is subsequently further explored.

Figure 5 shows per-cell precipitation trends across three initial calcium concentrations for simultaneous growth/mineralization trials. These values of per-cell precipitation are calculated similarly to those presented in Figure 3 and Figure S3, with the exception that the “final” calcium concentrations here are measured at the time of growth stoppage, rather than the time of post-processing (36 h). Data are presented in this way in order to specifically highlight how the population-averaged extent of encapsulation changes over time during the active growth period. Trends in per-cell precipitation over time also provide an important validation endpoint in subsequent modeling exercises (section 2.2.2).

**Figure 5.**
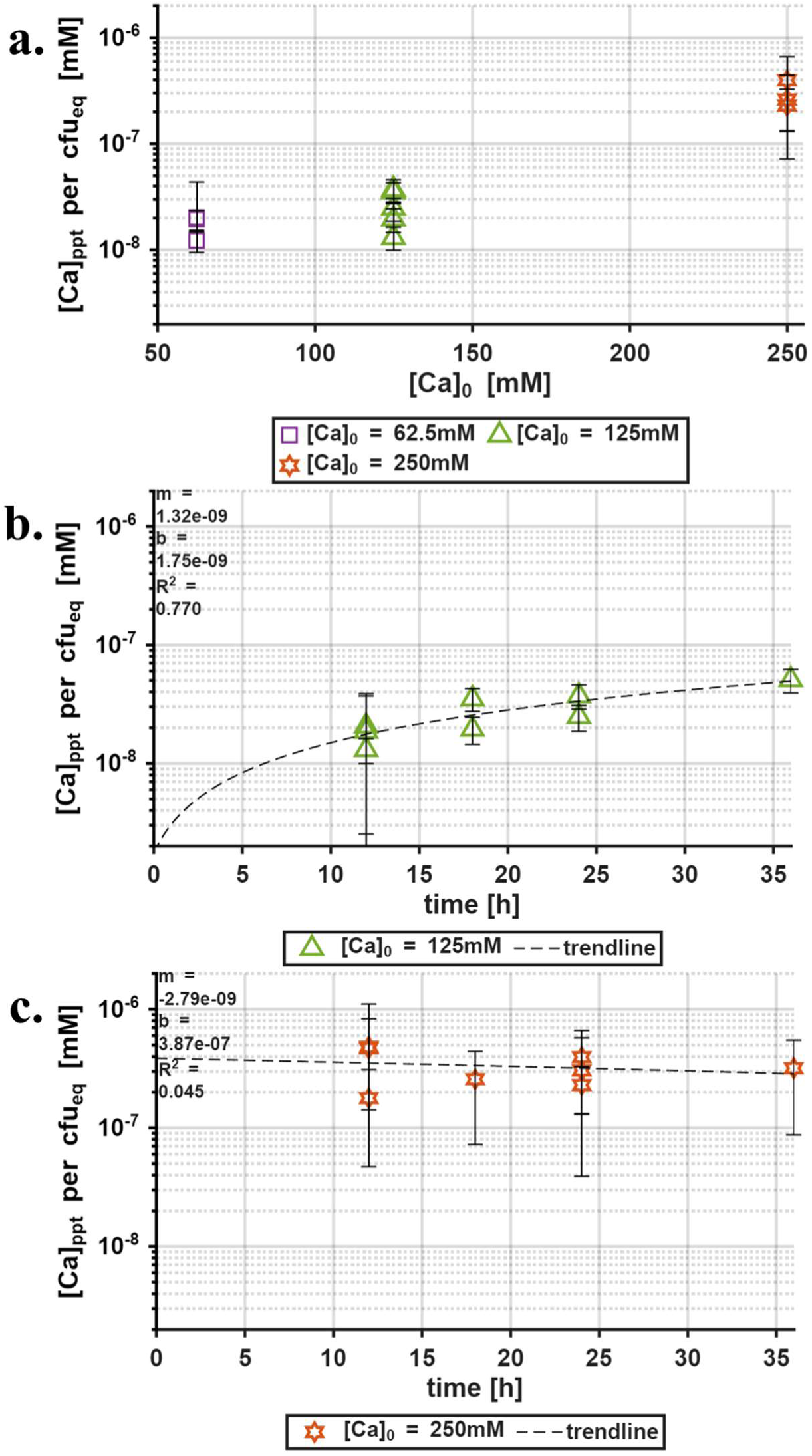
Estimates of per-cell precipitate accumulation in cultures with different initial calcium concentrations. Dissolved calcium measurements used in these estimates were taken at the time of growth stoppage, which is reported on the x-axis in (b) and (c). (a) shows the range of observed per-cell precipitation values in cultures of each initial calcium concentration, while (b) and (c) show the relationship between per-cell precipitation and culture growth time for cultures with initial calcium concentrations of 125 mM and 250 mM, respectively. Time-resolved data for 62.5 mM cultures are not shown here, as dissolved calcium generally became fully depleted in these cultures within 18 h, leaving too few datapoints to fit a trendline. As was done for Figure 3, a version of this figure was also prepared without screening samples for dissolved calcium depletion (Figure S7).

In samples with an initial calcium concentration of 250 mM, all per-cell precipitation values are in the range observed during demonstration trials (Figure 3, Figure S3) at all timepoints, while other initial calcium concentrations yield per-cell precipitation values one order of magnitude lower (Figure 5a). Although the underestimation of cell numbers at low cfu loadings (Figure 2, Figure S1) could inflate per-cell precipitation values in 250 mM samples, the magnitude of this effect is not large enough to explain even 50% of the observed difference. Examining trends in per-cell precipitation over time (Figure 5b-c) additionally shows that while encapsulation is, as expected, completed in the first 12 h for 250 mM samples, 125 mM samples show a linear increase in average per-cell precipitation over time. In the 36 h experimental timeframe, these populations never approach the full extent of encapsulation derived for this cell number range in section 2.1.2 (Figure 3, Figure S3).

Both the overall decrease in per-cell precipitation and its increasing trend over time are expected in actively growing cells, as new biomass is constantly being added for the first 18-24 h and precipitate accumulation necessarily lags growth. The size of this lag is, however, larger than would be expected in a primarily reaction-controlled environment, as previously shown from Figure 4. Furthermore, the population-wide rate of precipitate accumulation appears to be sensitive to calcium concentration. For example, while 125 mM cultures reached stationary phase in 12-24 h across all three growth/mineralization trials, precipitate accumulation continued until 36 h (Figure 5b) and remained almost 50% lower than the expected maximum (Figure 3, Figure S3). This is in contrast to the apparent complete encapsulation of cells in 250 mM samples within 12 h, noted above.

In summary, the cumulative evidence from trials on both inactive and growing cells suggests a nonlinear relationship between initial calcium concentration and cell growth, which may stem from mass transport constraints slowing precipitation to a degree inversely related calcium concentration. Successful coexistence of cell growth with mineralization in 62.5 and 125 mM samples appears to depend on growth rates sufficiently outpacing encapsulation rates at early times, when calcium is most abundant, whereas higher cell numbers and lower calcium concentrations at later times prevent most cells from ever becoming fully encapsulated. In contrast, it is hypothesized that under these experimental conditions, cells grown at 250 mM of dissolved calcium become inactivated through encapsulation before they are able to divide. As concentrations of 250 mM and higher are common in MICP studies, this implies that a robust understanding of realistic precipitation rates, likely a function of condition-specific mass transport constraints, is important to predicting the trajectory of systems designed for simultaneous growth and mineralization. The hypothesized relationship between mass transport, precipitation rate, and cell growth is further explored in two modeling exercises.

#### 2.2.2 Modeling the Relationship Between Encapsulation and Growth

The apparently nonlinear relationship between calcium concentration and cell growth, and its hypothesized relationship to encapsulation rate, motivated the modeling exercise done here. Model development was made feasible by the quantification of maximum per-cell precipitation during demonstration trials (Figure 3) and its validation in growth/mineralization trials (Figure 5). Models aimed to replicate the observed trends over time in cell growth and precipitation (Figure 4) as well as per-cell precipitation (Figure 5) in two ways, detailed in section 3.8. Although both models readily reproduced the precipitation behavior observed in trials on inactivated cells, an important difference emerged when simultaneous growth and precipitation was simulated.

Figure 6 compares experimental and modeled trends in growth and precipitation over time across the three initial calcium concentrations tested. While the diffusion-limited model appears to slightly underestimate precipitation relative to the completely-mixed model at initial calcium concentrations of 62.5 and 125 mM (Figure 6b, d), it reproduces the key observation that both growth and precipitation behavior change dramatically between 125 and 250 mM (Figure 6f), with almost negligible cell growth at the higher calcium concentration. The completely-mixed model was not able to reproduce this trend at any value of IT (Figure 6a, c, e; Figure S8). The diffusion-limited model was additionally capable of reproducing the differing trajectories of per-cell precipitation over time for initial calcium concentrations of 125 mM and 250 mM (Figure 5b-c), although actual values were underestimated in both cases (Figure S9). Because IT and RF_min_ were fit in part to reproduce the observed cell growth and calcium concentration trajectories, the diffusion-limited model’s ability to also capture the differing per-cell precipitation trajectories across calcium concentrations is not a fully independent validation. It is, however, a nontrivial additional check, since accurately fitting two trajectories individually does not guarantee that their ratio’s shape will also be reproduced.

**Figure 6.**
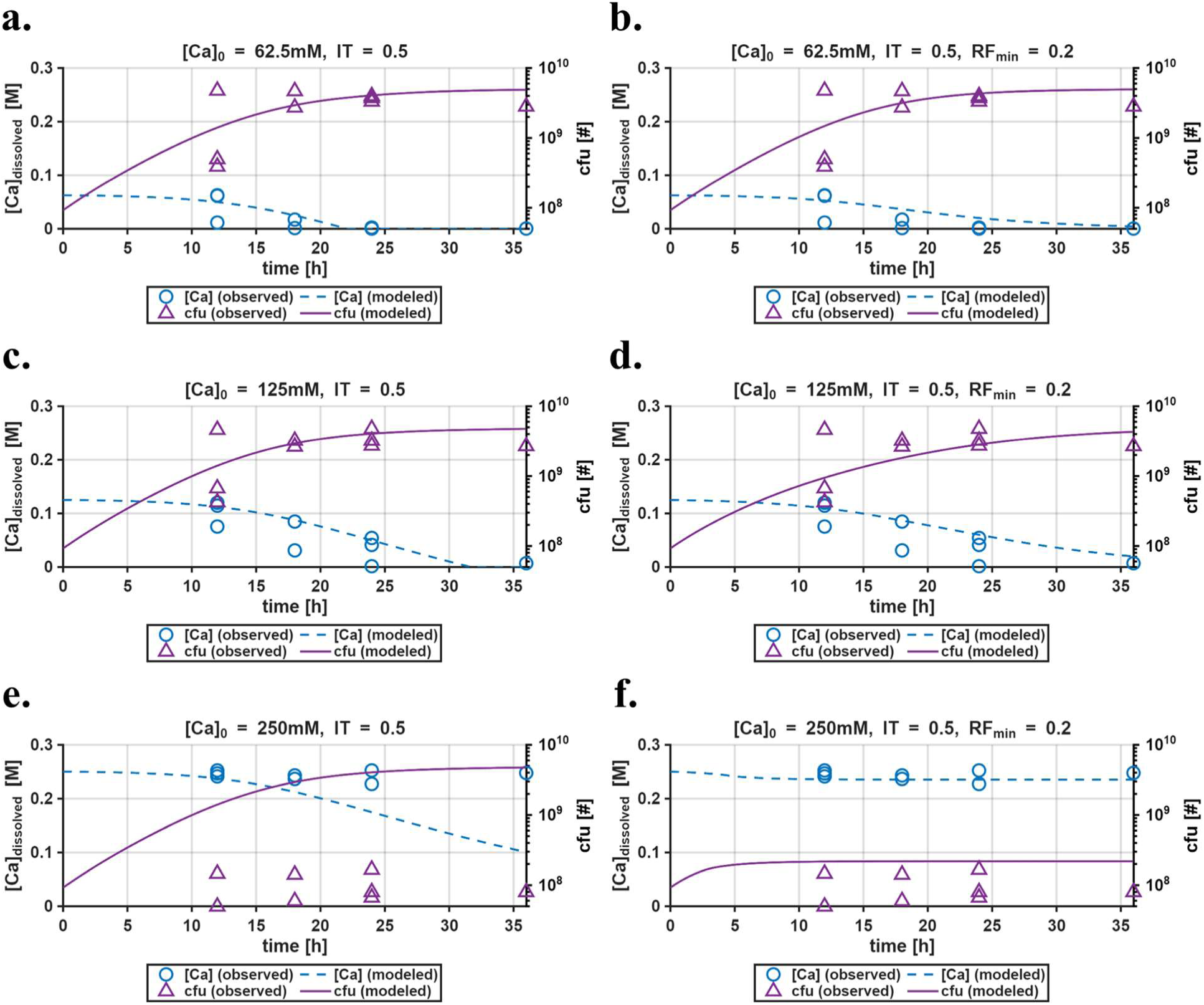
Cell number and carbonate precipitation trajectories predicted by two encapsulation models, for a scenario involving cell growth with initial calcium concentrations of 62.5 mM (a and b), 125 mM (c and d), and 250 mM (e and f).

Rather than attempting to fully describe the complex bio-physico-chemical dynamics underlying simultaneous microbial growth/precipitation processes, the completely-mixed and diffusion-limited models are intended to explore two possible sources of nonlinearity in the relationship between calcium concentration and cell growth. It was hypothesized in section 2.2.1 that the distinction between low and high calcium conditions is the amount of time required for precipitates to accumulate to the critical threshold at which a cell ceases to divide, here captured by the fitting parameter, IT. Based on the completely-mixed model’s poor fit, it appears that the additional time required to reach calcium carbonate saturation at lower initial dissolved calcium concentrations is not sufficient to explain the difference in growth behavior. In contrast, the diffusion-limited model’s improved fit shows that constructing a scaling function from mass transport parameters may allow models to better capture salient trends in cell growth across calcium concentrations, as well as the rate of growth in average per-cell precipitation over time. This suggests that parameterizing the mass transport constraints inherent to different physicochemical environments may be a useful starting point for capturing the influence of encapsulation in simultaneous growth/mineralization processes.

### 2.3 Implications and Future Work

By reporting a modified method for microbial biomass quantification and applying this method to simultaneous growth/mineralization processes, this study yields several insights that may support a more fundamental understanding of encapsulation in microbial mineralization systems. These findings also carry practical implications for engineered applications of microbial mineralization. For example, if the observed coupling between growth and encapsulation is found to generalize beyond the mixed batch system used here, it suggests that MICP injection or calcium dosing strategies could be deliberately designed to favor different balances of total precipitate accumulation, precipitation efficiency, and treatment speed, depending on the goals of a given application. Applying the tools developed here to these and related questions may eventually support more predictable and tunable outcomes in engineered microbial mineralization processes.

Accordingly, a number of questions remain for further investigation. On the methodological side, possible origins of the noted variation in DNA yield with both cell number and calcium concentration are outlined in section 2.1.1 but not explicitly interrogated here. Additionally, while only genomic DNA extraction was attempted here, exploring the efficacy of this method for mRNA extraction could substantially expand the method’s utility. Given the nature of most engineered microbial mineralization research, the method will also likely require further optimization in systems that include rock and/or soil matter to which both cells and nucleic acids may sorb.

Regarding the fundamental mechanisms governing simultaneous growth/mineralization processes, the biological mechanism responsible for growth suppression observed at 250 mM calcium is not fully resolved here. For example, it is possible that 250 mM of calcium induces stress independent of either cell encapsulation or the accounted-for ionic strength effect (section 3.1), and that this stress causes *S. pasteurii* to enter a dormant or sporulating state. Likewise, fully untangling the proposed encapsulation-induced growth suppression framework will require experiments at finer time resolution and more tightly-controlled mass transport regimes, in order to more conclusively assess the respective rates of growth and encapsulation and their relationship to mass transport constraints.

Similarly, the diffusion-limited model’s construction, used here as a tractable scaling device rather than a literal description of transport (section 3.8), does not by itself identify the physical origin of the local mass transport constraint it approximates, leaving open the roles of cell aggregation, cell crowding, and incomplete mixing, to name some examples. The potential importance of aggregation or floc formation is specifically indicated by trends in per-cell precipitation with cell number (section 2.1.2). More broadly, precipitate accumulation on cells is known to be spatially heterogeneous. Differences in crystal size, morphology, and polymorph coincide with differences in precipitation rate and EPS production^46,47^, and dissolution and reprecipitation processes are known to consolidate carbonate precipitates over timescales of hours^48–50^. All of these factors could influence the ease of mass transport around a partially encapsulated cell. While systematic investigation of these open questions would be valuable in terms of fundamental mechanistic insight into encapsulation, it is also important to note that realistic incorporation of mass transport limitations into models of microbial mineralization is substantially more complex under conditions of fluid flow through heterogeneous porous media. Further parameterizing encapsulation and transport constraints as a function of variables such as flow regime, particle surface chemistry, and biomass distribution will therefore be an essential step toward extending fundamental principles to field-relevant models.

## 3. Methodology

### 3.1 Medium Formulations and Cell Culture

Unless otherwise noted, all media used for cell growth contained a final concentration of 3g/L Difco Nutrient Broth, 0.15M ammonium chloride, and 0.30M urea, meant to be representative of commonly-used growth medium recipes^25,51^. 5x concentrated stock solutions of growth medium were prepared to facilitate the preparation of calcium-containing growth medium for experiments, which required initial calcium concentrations ranging from 0 to 250 mM. The desired final calcium concentration was achieved by adding different amounts of 1.250 M calcium chloride stock solution. To avoid confounding results with a range of different ionic strengths, a 3.750 M stock solution of sodium chloride was prepared and used to adjust ionic strength in samples that received less calcium, so that the initial ionic strength was a constant 0.750 M across all cultures. All solutions were either autoclaved (salt solutions) or filter-sterilized into autoclaved bottles (growth medium) prior to use. Hard agar streak plates used for colony isolation and storage were prepared by sterile vacuum filtration of 2x concentrated growth medium into 2x concentrated agar solution, both at 45°C, for a final concentration of 1.5% agar. Soft agar plates used in pour-plate colony enumeration assays were prepared similarly with a final concentration of 0.3% agar.

The MICP model organism *Sporosarcina pasteurii* (ATCC 11859) was used in this study. Prior to beginning experiments, cultures were inoculated from a single colony on a streak plate stored at 4°C, and grown for 24 h in 250 mL foam-stoppered flasks containing 100 mL of growth medium at 30°C and 120 rpm shaking. Fresh cultures of the same volume were inoculated with 0.5 mL of the overnight culture and grown for 12 h prior to beginning the experiment, in order to obtain cells in exponential growth phase. Following culture to exponential growth phase and prior to inoculation of experimental reaction tubes, growth medium was removed from cells by centrifuging at 4500 rcf for 10 min at 20°C, decanting, and resuspending the cell pellet in sterile isotonic saline solution (0.9% sodium chloride). This process was repeated twice before a final resuspension in saline solution. The optical density (600 nm, 1 cm path length) of cell suspension used for inoculation was adjusted to between 0.49 and 0.51. In order to obtain an approximate conversion between OD600 and total intact cell numbers, colony-forming units (cfu) per mL in the prepared suspension were estimated through colony count assays conducted in triplicate on soft (i.e. 0.3%) agar plates. Although cfu does not exactly correspond to total intact cells, it is assumed to be a close approximation in a fresh suspension of exponential-phase cells, and so cfu is also used as a proxy for total intact cell numbers.

### 3.2 Demonstration Trials Using Inactive *S. pasteurii*

#### 3.2.1 Demineralization Method Validation

Assessing the efficacy of the proposed demineralization method required using known quantities of whole cells to carry out carbonate precipitation in growth/mineralization medium. Sodium azide was therefore added to the growth/mineralization medium in order to prevent growth during demonstration trials. Preliminary experiments to determine a suitable concentration of sodium azide are documented in the supplementary information (Table S2 and Figure S10), along with preliminary experiments used to estimate approximate timescales of mineralization and demineralization and identify the optimal EDTA concentration for the demineralization solution.

Mineralization reactions were prepared in 50 mL Eppendorf DNA LoBind centrifuge tubes. The total volume of growth/mineralization medium and cell inoculum added to reaction tubes was 20 mL, and 2 mL of 5.5% sodium azide stock was added to this for a final volume of 22 mL and final concentration of 0.5%. After preparing reaction tubes, tubes were capped, inverted to mix, and placed in a 30°C incubator at 180 rpm shaking for 36 h.

As precipitates often adhered to tube walls, demineralization was carried out in reaction tubes. Precipitates and cells were separated from the mineralization medium by centrifugation at 4500 rcf for 30 min, after which the mineralization medium was decanted and replaced with a demineralization solution consisting of 0.25 M EDTA at a pH of 8.0, as well as 0.10 M Tris at pH 8.0 (important for matching pH to EDTA-negative “non-demineralized” control) and 0.05% sodium azide. Since the highest concentration of calcium used was 0.25 M, adding a volume of demineralization solution equal to the mineralization medium volume (20 mL) was theoretically sufficient to demineralize all precipitates that might have formed. Demineralization occurred over 24 h, also at 30°C and 180 rpm. Subsequently, cells and cell debris were separated from the demineralization supernatant by centrifugation for 30 min at 4500 rcf, and each component was subjected to a separate DNA extraction protocol.

Cell pellets were resuspended in 1.5 mL dilute Tris/EDTA (10 mM/0.5 mM), transferred to 2 mL Eppendorf DNA LoBind microcentrifuge tubes, and spun at 10000 rcf for 2 min. This was repeated once in order to rinse cells of demineralization medium, after which DNA extractions were performed using Qiagen UltraClean Microbial DNA kits according to the manufacturer’s instructions.

DNA was concentrated from the demineralization supernatant by adding 4 mL of 6x concentrated sodium citrate (pH 7.5) for a final concentration of 0.30 M and final volume of 24 mL. 1.2 mL of this solution was added to 800 μL of molecular biology-grade isopropanol in an Eppendorf DNA LoBind 2 mL microcentrifuge tube at room temperature. After inverting to mix, DNA was pelleted in Eppendorf DNA LoBind 2 mL microcentrifuge tubes at 12500 rcf for 15 min at 4°C, and supernatant was decanted. This was repeated five times for each sample so that a total of 6mL was concentrated, representing 25% of the total DNA in the demineralization supernatant. Two rinses were done with 70% ethanol in order to de-salt the DNA pellet, and pellets were resuspended in 200 μL Tris/EDTA buffer after air drying for 45 min.

Three of the above-described demonstration trials were carried out. Each trial consisted of twelve samples across four calcium concentrations (0, 62.5, 125, and 250 mM) and three cfu loadings (7.0x10^7^, 2.8x10^8^, and 1.1x10^9^ total cfu; these are equivalent to 3.5x10^6^, 1.4x10^7^, and 5.5x10^8^ cfu/mL).

#### 3.2.2 Negative Controls

Two sets of control experiments were performed in order to provide a basis for comparing DNA yields. In the first set of control experiments, termed the “no demineralization” control, mineralization reactions were carried out as described above using 1.1x10^9^ total cfu and 250 mM calcium, and the Qiagen kit was used to extract DNA directly from the resulting precipitates. In the second, termed the “no mineralization” control, the Qiagen kit was used to extract DNA from cell pellets of 2.8x10^8^ total cfu. Extractions were performed immediately after cell growth to exponential phase and saline rinse (section 3.1), so cells were never diluted into the 20 mL reaction tubes used in other trials. Colony count assays were also performed on cell suspensions used in “no mineralization” controls, in order to establish the expected value and variance in DNA yield per cfu (Table S1). DNA yield was quantified using real-time PCR, described below (section 3.5).

### 3.3 Simultaneous Growth/Mineralization Trials

Three growth trials were performed with the same conditions and medium compositions as demonstration trials but without the addition of sodium azide. 7.0x10^7^ cfu were used for initial inoculation, and four sets of tubes were prepared for each of the four calcium concentrations tested (0, 62.5, 125, and 250 mM). Since preliminary data indicated that the onset of exponential growth may not occur until 8-10 h after inoculation, the first trial used timepoints of 12, 24, and 36 h. At each timepoint, one tube of each calcium concentration was retrieved and sodium azide was added to a final concentration of 0.5% in order to stop cell growth. Just prior to adding sodium azide, additional characterization of dissolved oxygen, pH, and dissolved calcium concentration was also completed, as described below. Since no significant growth was found to occur after 24 h, the second and third growth trials used timepoints of 12, 18, and 24 h. Demineralization and DNA extraction were initiated at the same time for all samples, 36 h after initial inoculation, following the same procedure used in demonstration trials.

### 3.4 Dissolved Calcium, Dissolved Oxygen, and pH Measurements

Dissolved calcium concentrations were measured by inductively-coupled plasma mass spectrometry (ICP/MS). 150 μL samples were taken from reaction tubes and diluted into 2% nitric acid prior to analysis, with a final volume of 15 mL. In both demonstration and growth trials, ICP/MS samples were taken at the start of reactions and again at 36 h, just before initiating demineralization. In growth trials, ICP/MS samples were also taken alongside pH and dissolved oxygen measurements just prior to adding sodium azide to stop growth. In order to characterize the culture medium chemistry as accurately as possible (i.e., avoid capturing precipitated calcium in ICP/MS measurements and minimize re-equilibration with atmospheric air before electrode measurements), ICP/MS sampling and electrode measurements were preceded by centrifuging tubes without opening for 20 min at 4500 rcf and 20°C. After removing caps, ICP/MS samples were taken first, followed by dissolved oxygen and pH measurements, if applicable.

### 3.5 DNA Quantification with Quantitative PCR (qPCR)

DNA samples were stored at -20°C for up to three weeks prior to qPCR quantification. Primers (F: GGCTTGAGTACAGAAGAGGAAAG; R: GCGTCAGTTACAGACCAGAAAG) were developed for a 141-bp amplicon in the 16S rRNA V3-V4 region of *S. pasteurii*^39^. Since experiments began with pure cultures of *S. pasteurii* and aseptic techniques were observed throughout, primer specificity to *S. pasteurii* was deemed nonessential. Nonetheless, BLAST analysis was carried out to identify other targets that might be amplified. In addition to *Sporosarcina*, the forward primer sequence was also found in the *Paenibacillus* genus, while the reverse primer sequence appears to be highly conserved across the Bacillaceae family. Primers and corresponding gBlock (for standard curve) were ordered from Integrated DNA Technologies. For qPCR, triplicate reactions of 12μL volume each were prepared using Fisher Scientific SYBR Green qPCR master mix, on Applied Biosystems MicroAmp Fast Optical 96-well 0.1mL plates. Amplification was performed on a Thermo Fisher QuantStudio 3 PCR system according to the master mix manufacturer’s suggested cycling conditions for normal-speed amplification. Standard curves for each amplification are shown in Figure S11. DNA yields are reported as the total number of gene copies (gc) recovered for each sample.

### 3.6 Estimating Cfu_eq_ and Per-Cell Precipitation

From raw data on dissolved calcium concentration (section 3.4) and gc recovered (section 3.5), two additional quantities are calculated in order to simplify analyses. Cfu equivalents (cfu_eq_) are defined as the number of gc recovered from a sample divided by the number of gc recovered from a reference sample and multiplied by the estimated number of cfus (itself a proxy for intact cell numbers, as described in section 3.1) in the reference sample. Here, reference sample properties are taken to be the average of the “no mineralization” control trials (section 3.2.2, Table S1), as further explained in section 2.1.1. Standard deviations of each estimate are propagated into the final cfu_eq_ value as described in section 3.7. As “no mineralization” control trials used suspensions of fresh, exponential-phase cells, cfu numbers are expected to be a suitable proxy for the total number of intact cells in these samples, as discussed in section 3.1. Cfu_eq_ is therefore adopted as a convenient means of estimating the number of intact cells in a sample from the number of gene copies recovered. In other words, the term “cfu_eq_” reflects the calibration method used to derive this quantity, not an expectation that it will be equal to the actual number of culturable cells in a sample.

Second, the per-cell precipitation is defined as the difference between the initial and final dissolved calcium concentrations, divided by either the number of intact cells present (if known) or its gc-based estimate, cfu_eq_. In demonstration trials on inactive cells, the two methods of calculating per-cell precipitation are compared in order to assess the reliability of cfu_eq_ as an estimator. The primary motivation for quantifying per-cell precipitation is to establish a benchmark level of precipitate accumulation that represents full encapsulation (i.e., the point at which precipitation cannot continue even with excess dissolved calcium), against which partially-mineralized samples can be compared in order to assess cells’ average degree of encapsulation. This benchmark value, which is expected to be a function of variables such as medium chemistry and mixing, is derived for these experimental conditions using data from demonstration trials on inactive cells. It is then used in the analysis of growth/mineralization trials (section 3.3) and as a parameter in the simplified encapsulation models developed here (section 3.8). Note that the per-cell precipitation represents a population-wide average for all cells in a sample, and is therefore sensitive to processes such as aggregation and flocculation. Particular caution must be applied to analyses of growing samples, as different degrees of encapsulation are expected among differently-aged cells even in the absence of aggregation. Unless otherwise specified, per-cell precipitation is only calculated for samples whose final dissolved calcium concentration was between 10% and 90% of the initial value. These screening criteria select for samples where significant precipitation occurred and where dissolved calcium was not fully depleted, both of which are necessary conditions for per-cell precipitation to be meaningful.

### 3.7 Statistical Analyses

Error bars shown in figures and ranges reported on numerical values represent standard deviations (i.e. the square root of the variance), where the datapoint or numerical value itself is the mean. Unless otherwise specified, standard deviations are obtained from triplicate qPCR, ICP/MS, or colony count results. In many cases, it is informative to report quantities that are the products of two or more means, each with their own standard deviation, such as cfu_eq_ (defined in section 3.6). When this is done, error was propagated using standard techniques^52^. When statistical significance is assessed for the difference between two sample means, Welch’s t test is applied with the assumption of unequal variance. Finally, all linear trendlines are fit using ordinary least squares regression (OLS). In order to assess whether differences in trendline slope or y-intercept are statistically significant, confidence intervals are constructed assuming a Student’s t distribution, with confidence interval width selected so that overlapping confidence intervals indicate lack of a statistically significant difference at the desired p-value^53,54^.

### 3.8 Modeling

In an effort to better understand the relationship between encapsulation and calcium concentration, experimental data were used to develop two simplified models of simultaneous cell growth and precipitation. Figure S12 graphically depicts the key concepts underpinning these models, while Figure S13 summarizes the constants and governing equations used in each. In both models, cells are assumed to be homogeneously distributed throughout the bulk solution. The models’ approach to growth, ureolysis, and precipitation activity involves sorting cells into age cohorts according to the time step in which they were formed (Figure S12a). As a given cohort ages, calcium carbonate precipitates are assumed to accumulate on all cells in the cohort at the same rate. While the rate of precipitate accumulation on cells is always derived from whole cell ureolysis kinetics^45^, the core difference between the two models is how raw ureolysis rates are converted to precipitation rates, discussed below.

In both models, precipitate accumulation on cells causes the cells’ ureolysis rate to decline linearly, reaching zero when per-cell precipitation reaches the empirically-estimated maximum (section 3.6). Growth rates decline in a similar manner to ureolysis rates with the addition of a fitting parameter termed the inactivation threshold (“IT”), the theoretical level of precipitate accumulation at which cells cease to divide. As previously shown^51,55–57^ and confirmed here in experiments on inactive cells (section 2.1.1), ureolysis activity is largely unrelated to cell viability or growth conditions. Therefore, both models assume that the extent of encapsulation controls the rates of both ureolysis and growth independently of one another, with each rate being expressed as a fraction of its maximum value (Figure S13). Cell inactivation will intuitively occur at a lower degree of encapsulation than total isolation from the environment (i.e. the point at which ureolysis activity reaches zero), which is captured here in the inactivation threshold fitting parameter, IT.

The first model, named the “completely-mixed” model, determines the amount of precipitation at each time step by using the total number of ureolysis reactions to iteratively solve for the equilibrium concentrations of all relevant chemical species, assuming perfect mixing. This is in contrast to single-reaction approaches often employed in MICP models, which are reported to be as effective as equilibrium modeling while being less computationally costly^24,58^. In this case, the motivation behind equilibrium modeling was to attempt to simulate a realistic time delay between ureolysis and precipitation, as precipitation will only occur once calcium carbonate saturation has been achieved.

The second model, named the “diffusion-limited” model, adopts a single-reaction approach to precipitation, which usually implies carbonate precipitation in a one-to-one ratio with ureolysis. Here, however, a scaling function is used to adjust the rate of precipitation downward from the ureolysis rate, with the size of the adjustment inversely proportional to the calcium concentration. To obtain a scaling function, two reference volumes are calculated for each cell (Figure S12b). The reaction reference volume, V_rxn_, represents the volume of solution that would be depleted of calcium in one time step if precipitation proceeded in a one-to-one ratio with ureolysis. This volume is re-calculated each timestep as calcium concentration changes. The diffusion-limited reference volume, V_diffn_, represents the volume enclosed by a sphere centered on the cell with a radius equal to the characteristic diffusion length scale for one time step. This volume is calculated once, remaining constant in time. The ratio V_diffn_/V_rxn_ is then used to scale the cell’s precipitation rate down from the ureolysis rate. A second fitting parameter, RF_min_, is adopted in order to prevent the scaling factor from approaching zero at low but nonzero calcium concentrations.

Physically, the diffusion-limited model construction represents a limiting-case scenario in which local mass transport occurs only through molecular diffusion, but this is not intended as a literal description of transport in the mixed reactors used in this study. Rather, this construction is used to generate a smooth, calcium concentration-dependent scaling for precipitation rate, whose overall form (with the exception of the RF_min_ parameter) is derived from independent physical and kinetic parameters rather than fit directly to empirical data.

## Acknowledgements

The authors would like to thank Prof. Krista Wigginton and her research group, particularly Dr. Ke Zhang, Abigail Monahan, and Allyson Criswell, for their support with planning, training, and equipment access related to the qPCR work in this manuscript. The authors also thank Dr. Peng Peng for performing preliminary PCR tests during method development. This work was made possible by the financial support of the Rackham Graduate School at the University of Michigan, through the Rackham Graduate Student Research Grant and Rackham Predoctoral Fellowship programs.

